# Microbial Treatment of Soil Contaminated with Spent engine Oil / Biotreatment of Soil Contaminated with Spent Engine by Microorganisms

**DOI:** 10.1101/268185

**Authors:** A. A. Ayandele

## Abstract

The potential of six microorganisms (*Pseudomonas aeruginosa, Micrococcus* sp, *Flavobacterium* sp, *Rhizopus* sp, *Penicillium* sp and *Fusarium* sp) isolated from hydrocarbon contaminated site were evaluated for their biodegradation ability. The soil samples were contaminated with 5% (w/v) of spent engine oil and the rate of biodegradation of the oil was studied for a period of 10weeks under greenhouse experiment. The total heterotrophic bacteria count (THBC), total hydrocarbon degrading bacteria count (THDBC), physicochemical and heavy metals properties of the soil samples and Total Petroleum Hydrocarbon (TPH) were determined after treatment with test organisms. THBC and THDBC ranged from 0.175 to 0.280 CFUg^-1^ and 0.47 CFUg^-1^ respectively for the control plot, while THBC is ranging from 0.197 to 0.275 CFUg^-1^ and THDBC was 0.180 to 0. 473 CFUg^-1^ for the contaminated plot. There was a slight increase in the pH value of the contaminated soil sample and the treated soil samples as the experimental weeks increased. The results obtained showed a significant decrease (at p ≤ 0.05) in the nutrients content of the soil samples. There was an increase from 1.09 in the control to 15.5% in the content of organic matter after contamination and from 1.88% to 26.8% in the % of organic matter too. There was a significant reduction (at p ≤ 0.05) in the concentration of Fe, Zn, Pb, Cd, Cu, Cr and Ni after 10 weeks of incubation with the tested organisms. Plant growth in the treated contaminated soil samples ranged from 32.6cm to 38.6cm, while that of the control 1 (Uncontaminated soil) was 51.2cm and 19.7cm high was observed in the Control 2 (contaminated untreated soil) after 22 days of the experiment. The TPH degradation (% loss) ranged from 79.7 to 89. 2% after 10 weeks of treatment. *P. aeruginosa* had the highest level of degradation (89.2%), while *Micrococcus* sp and *Rhizopus* sp had the least degradation at 79.9%.

All the microorganisms used in this study had the abilities to remediating soil contaminated with spent engine oil and the remediated soil samples were able to support the growth of *Zea mays* at 5% (w/v) level of contamination.

## INTRODUCTION

Engine oil could simply be defined as a thick mineral liquid applied to a machine or engine so as to reduce friction between the moving parts of the machine (Shahida *et al.*, 2015). Used engine oil as the name implies represent oil that has undergone destructive changes in the property when subjected to oxygen, combustion gases, and high temperature. The said oil also undergoes viscosity changes as well as additive depletion and oxidation (Mark *et al.*, 1982).

In Nigeria, it is common among motor mechanics to dispose of spent engine oil into gutters, water drains and soil (Okonokhua *et al.*, 2007). Spent engine oils contain high percentage of aromatic and aliphatic hydrocarbons, nitrogen, sulphur compounds, and metals (Zn, Pb, Cr and Fe) than fresh oils, some of these metals in used engine oil can dissolve in water and move through the soil easily and may be found in surface water and groundwater (Mohd *et al.*, 2011; Abdulsalam *et al.*, 2012). Spent engine oil causes great damage to soil and soil microflora. It creates an unsatisfactory condition for life in the soil due to poor aeration, immobilization of soil nutrients and lowering of soil pH (Ugoh and Moneke, 2011). It has been shown that marked changes occur in soil contaminated with hydrocarbons and these changes affect the physical, chemical and microbiological properties of the soil (Okonokhua *et al.*, 2007).

Microbial degradation is the major mechanism for the elimination of used petroleum products from the environment (Bartha, 1992). Soils contain very large numbers of microorganisms which can include a number of hydrocarbons utilizing bacteria and fungi (Namkoong *et al.*, 2002). Microorganisms are capable of breaking down many complex molecules by adaptation of their degradative enzyme system (Boonchan *et al.*, 2000). Bioremediation is, therefore, the application of naturally occurring process by which microorganisms transform environmental contaminants into harmless end products (Abdulsalam *et al.*, 2012).

Therefore, the objectives of this study were to determine the biodegradation ability of fungi and bacteria isolated from used engine oil contaminated soil, the effects of bioremediation on the physicochemical properties and heavy metals present in the contaminated soil.

## MATERIALS AND METHODS

### Field Work

Four experimental plots (1.0 by 1.0 m^2^) were mapped out for this experiment. Two plots were contaminated with 3L of spent engine oil and mixed thoroughly up to a depth of 10cm, while the remaining two plots which were not mixed with any spent engine oil served as a control. The soil samples were mixed together for aeration and watered every 3 days. After a week, Tze-e yellow maize seeds collected from International Institute of Tropical Agriculture (IITA), Ibadan, Nigeria, were planted on each plot. Maize plant heights were measured in cm and monitored for a period of one month. Determination of Total Oil-Degrading bacteria and Total Heterotrophic bacteria counts were also carried out every month for a period of nine months.

### Isolation of Microorganisms

#### Total Heterotrophic Bacteria (THB) Counts

A THB count was determined by pour plate method. Serial dilution was carried out on soil sample collected from each plot and 1ml of the aliquot from each of the dilution was inoculated by pour plate method onto Nutrient Agar plates in duplicates. The plates were incubated at 37°C. THB counts were determined after 24hrs of incubation.

#### Total Oil Degraders (TOD) Counts

Spent engine oil was collected from a mechanic shop and sterilized by using Millipore filter. 1ml of an aliquot of serially diluted soil sample was added onto Bushnell Hass agar containing 2ml of spent engine oil by spread plate method. The plates were incubated at 25°C and TOD counts were determined after 7 days of incubation.

#### Preliminary Screening of Spent Engine Oil Degraders

Nutrient and Potato Dextrose Agar were used for the isolation of bacteria and fungi respectively from contaminated soil samples. Pour plate method was also used as described above. Pure bacterial and fungal isolates were then streaked on oil agar medium to determine those isolates that can utilize spent engine oil as their sole source of carbon. The oil agar medium consisted of basal (Mineral Salts) medium; 1.8g K_2_HPO_4_, 1.02g KH_2_PO_4_, 4.0g NH_4_Cl, 2.0g MgSO_4_.7H_2_O, 0.1g NaCl, 0.1g Yeast extract, 0.05g FeCl_2_ and trace elements consisting of 0.1g H_3_BO_3_, 0.1g ZnSO_4_ and 0.4g MnSO_4_.H_2_O in 1L of sterile distilled water. The oil agar medium was divided into two; Nystatin was added to the medium used for bacteria to prevent fungal growth and Streptomycin to medium for fungal growth. Those isolates that showed good growth on oil agar medium were used for biodegradation experiment.

#### Characterization of Bacterial and Fungal Isolates

Microscopic and biochemical tests were used for determination of bacterial isolates and the isolates were characterized as described by Bergey’s Manual of Determinative Bacteriology (Holt *et al.*, 1994). While both macroscopic and microscopic analyses were carried out on fungal isolates to determine their identities according to Mackie and McCartney, (1999).

### Greenhouse Studies

#### Preparation of Seed Bags

Each seed bag contained 1.5kg of sterilized soil mixed with 150mL of spent engine oil at a ratio of 1:1w/v. Many seed bags were prepared for this experiment. Sterilized soil without spent engine oil was prepared to serve as positive control.

#### Preparation of Inocula Treatment and Seedling

Three bacterial and fungal isolates each were used for this experiment. Bacteria and fungi were inoculated into plates containing Nutrient and Potato Dextrose agar respectively for 48hrs. Isolates were centrifuged for 20 minutes, the supernatant was decanted, cells washed and resuspended in sterile saline water (0.9%). Bacterial and fungal cells were standardized by using Spectrophotometer. 100mL of each isolate was mixed separately with each seed bag containing soil sample and sterilized spent engine oil.

Seed bag containing sterile soil sample devoid of microorganisms served as positive control and was coded PC, Seed bag containing sterilized soil and spent engine oil but devoid of microorganisms served as negative control and coded NC,

Seed bag containing sterilized soil and spent engine oil with each of 1^st^, 2^nd^, and 3^rd^ bacterial isolates were coded BS1, BS2, and BS3 respectively,

Seed bag containing sterilized soil and spent engine oil with each of 1^st^, 2^nd^, and 3^rd^ fungal isolates were coded FS1, FS2, and FS3 respectively. All the treated soil samples and both the positive and negative control soil samples were duplicated.

Changes in Total Petroleum Hydrocarbons levels, physicochemical and heavy metal analyses were determined during the 10 weeks of the experiment.

#### Planting of Maize Seeds

Two viable seeds of Tze-e yellow maize were planted in each of the seed bags. Germination of maize seeds was monitored and plant heights were measured in cm and recorded for a period of two weeks. Watering of seed bags was carried out at every 5 days interval.

#### Physicochemical Analyses

Soil pH was determined by using a pH meter. Calcium and Magnesium were determined by titrimetric method (EL Mahi *et al.*, 1987). Total Nitrogen was determined by Kiehl digestion and steam distillation method (Sankaram, 1996). Available Phosphorus was also determined according to the method of Olsen *et al.* (1954), while Potassium was also determined by using the Flame photometer (Sankaram, 1996).

#### Determination of Total Organic Carbon and Organic Matter

The soil sample is sieved through 1mm sieve and 1g of sieved soil sample was put in a 100mL flask. 10mL potassium dichromate and 20mL sulphuric acid were added to the sample and shake very well. The mixture is allowed to cool on asbestos sheet and the volume is made up of 100mLwith distilled water and kept overnight. The optical density was measured at 660nm wavelength using a spectrophotometer.

% Organic Carbon = optical density × Factor F

% Organic Matter is determined by = Organic Carbon × 1.724

#### Determination of Total Petroleum Hydrocarbon (TPH)

Total petroleum hydrocarbon of the soil sample was determined by the method of Intergovernmental Oceanographic Commission (IOC) as described by Chukwujindu *et al.* (2008).

#### Heavy Metal Analysis

The heavy metal analysis was carried by Atomic Absorption Spectrophotometry (AAS). For each of the metal [Iron (Fe), Zinc (Zn), Lead (Pb), Cadmium (Cd), Copper (Cu), Chromium (Cr) and Nickel (Ni)], AAS is calibrated using a standard solution of the metal. 5g of soil sample was digested in 20mL of hydrogen chloride on a heating mantle to dryness. The extract was aspirated directly into the AAS and reagent blank was used to estimate the amount of each metal.

**Statistical Analysis**: Mean values of all the parameters taken were calculated and factorial analysis of variance (ANOVA) was also determined at 0.05 of the confidence interval.

## RESULTS AND DISCUSSION

Figures 1 and 2 showed the plant heights of maize planted on the contaminated soil and seed bags. Germination of maize seeds was delayed for the maize planted on the contaminated soil while there was no delay on the maize seeds planted on control plot, which is soil without spent engine oil (Fig 1). On day 6, maize plant height on control plot was 8cm while no visible growth was observed on a contaminated plot. By the 28^th^ days, plant heights on control and contaminated plots had increased to 88 cm and 42 cm respectively. Heights of maize planted inside seed bags containing spent engine oil were treated with different microorganisms to determine their effects on the growth of maize seeds, while control 1 (uncontaminated soil in seed bag) and Control 2 (uncontaminated soil seed bag) also contained maize seeds. While the maize plants showed varying heights on soil samples treated microorganisms, control 1 and control 2 (Fig 2). Control 1 had the highest plant height followed by soil treated with *Penicillin* sp, soil treated with *Rhizopus* sp showed the least maize plant height among the treated soil samples while control 2 showed the least plant height (Fig 2). Delayed in plant germination has been reported by Onwuka *et al.* (2012), while Adenipekun *et al.* (2013) has reported low leaves yield in *Corchorus olitorius* (L) growing in soil contaminated with spent diesel oil. It has been reported that hydrocarbon oil reduced soil quality and crop yield (Adeoye *et al.*, 2005).

**Figure 1:**
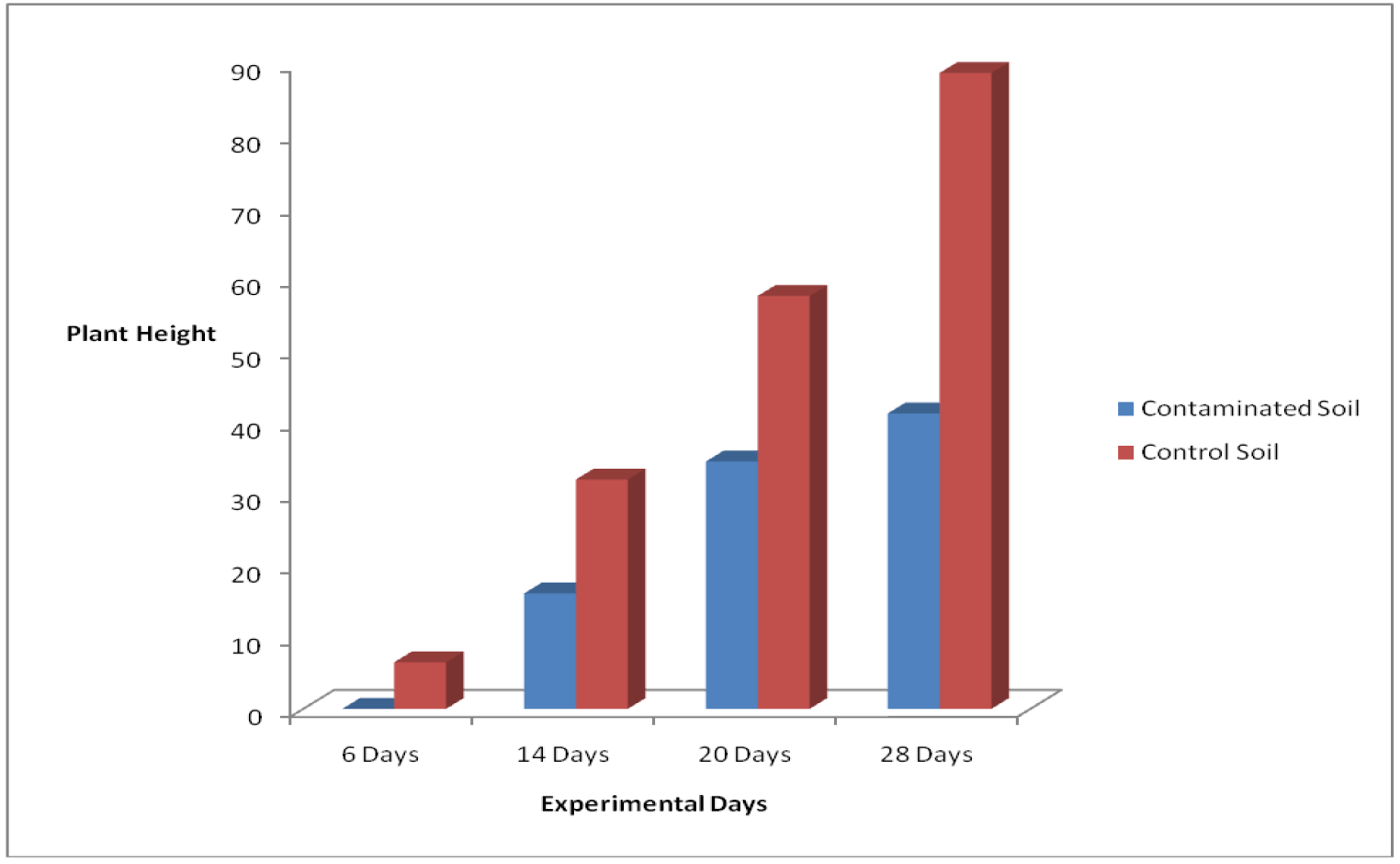
Maize Plant Height on Contaminated and Control Plots

**Fig 2:**
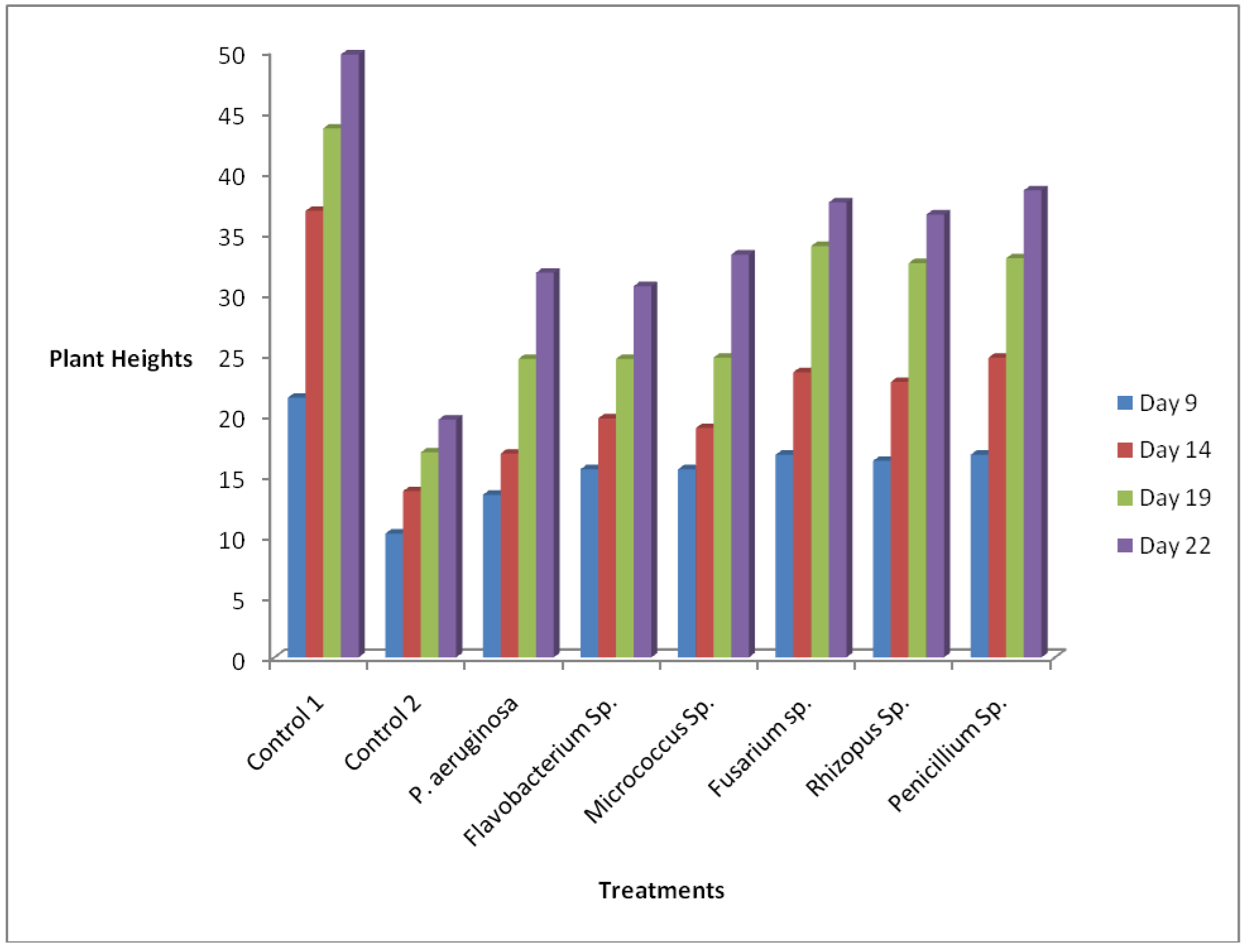
Maize Plant Heights with Different Treatment in Seed Bags

Isolation of both total heterotrophic and degraders counts was carried out monthly for nine months on both contaminated and control plots soil samples. Table 1 shows the counts of total heterotrophic and oil-degrading bacteria on both the uncontaminated (control) and contaminated plots. There was a gradual increase in the heterotrophic count on the control plot but as the experimental months increased, the counts started increasing from 248 CFU × 10^3^ in the first month to 280 CFU × 10^3^ (fifth month) but as the month progresses, it started reducing unto the last month but the counts of heterotrophic bacteria was higher in the contaminated soil samples at the beginning of the experiment (275 CFU × 10^3^) but as the month progressed, there was a reduction in the count but later increased to 297 CFU × 10^3^ on the last month. There was a decrease in the total oil degraders in the control plot from 95 CFU × 10^3^ to 47 CFU × 10^3^ in the control plot, while oil degraders increased from 180 CFU × 10^3^ to 473 CFU × 10^3^ on the contaminated plot by the fifth month but started reducing on the seventh month till it reduced to 325 CFU × 10^3^ on the last month of the experiment (Table 1). Subathra *et al.* (2013) also reported similar observation in their work. High heterotrophic bacteria counts in the contaminated plot towards the end of the experiment have been attributed to the fact that more nutrients are now available for the indigenous bacteria present in the soil due to the breakdown of hydrocarbons to the simpler forms by the oil degraders bacteria (Olubunmi, 2014). Rahman *et al.* (2002) also reported that population level of hydrocarbon utilizers and their populations within the microbial community appear to be a sensitive index of environmental exposure to hydrocarbons.

**Table 1:** Total Heterotrophic Bacteria and Total Hydrocarbon-Degrading Bacteria Counts in Control and Contaminated Soil Samples

| Sampling Months | Control Plot |  | Contaminated Plot |  |
| --- | --- | --- | --- | --- |
|  | Total Heterotrophic count | Total Oil Degraders Count | Total Heterotrophic count | Total Oil Degraders Count |
|  | ( CFU × 10<sup>3</sup>) | ( CFU × 10<sup>3</sup>) | ( CFU × 10<sup>3</sup>) | ( CFU × 10<sup>3</sup>) |
| June | 248 | 95 | 275 | 180 |
| August | 275 | 86 | 220 | 376 |
| October | 280 | 74 | 197 | 473 |
| December | 225 | 65 | 248 | 415 |
| February | 175 | 47 | 297 | 325 |

**Table 2:**
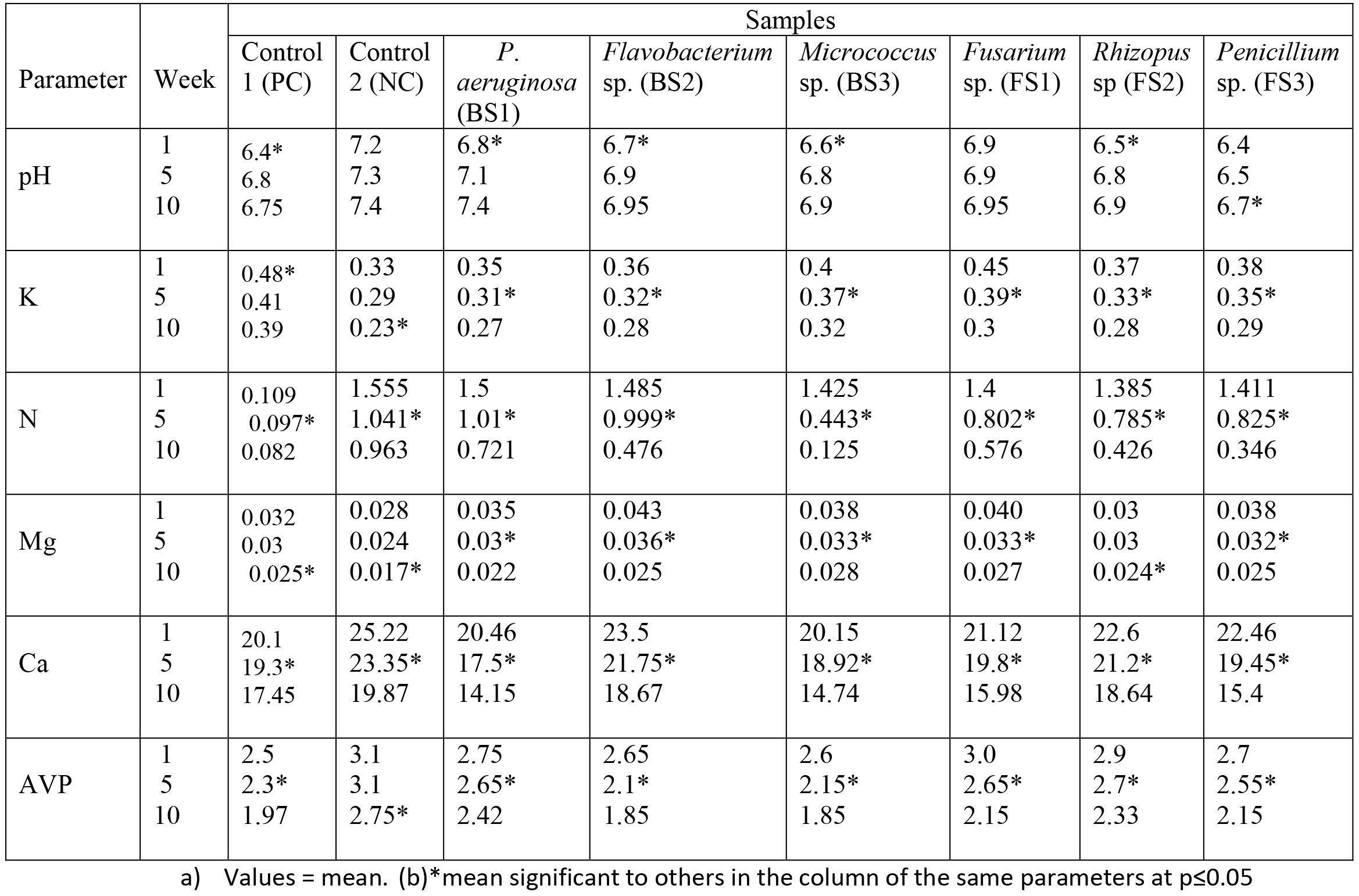
Physicochemical Analysis of Soil Samples with Different Treatments

| Parameter | Week | Samples |  |  |  |  |  |  |  |
| --- | --- | --- | --- | --- | --- | --- | --- | --- | --- |
|  |  | Control 1 (PC) | Control 2 (NC) | *P. aeruginosa* (BS1) | *Flavobacterium* sp. (BS2) | *Micrococcus* sp. (BS3) | *Fusarium* sp. (FS1) | *Rhizopus* sp (FS2) | *Penicillium* sp. (FS3) |
| pH | 1 | 6.4* | 7.2 | 6.8* | 6.7* | 6.6* | 6.9 | 6.5* | 6.4 |
|  | 5 | 6.8 | 7.3 | 7.1 | 6.9 | 6.8 | 6.9 | 6.8 | 6.5 |
|  | 10 | 6.75 | 7.4 | 7.4 | 6.95 | 6.9 | 6.95 | 6.9 | 6.7* |
| K | 1 | 0.48* | 0.33 | 0.35 | 0.36 | 0.4 | 0.45 | 0.37 | 0.38 |
|  | 5 | 0.41 | 0.29 | 0.31* | 0.32* | 0.37* | 0.39* | 0.33* | 0.35* |
|  | 10 | 0.39 | 0.23* | 0.27 | 0.28 | 0.32 | 0.3 | 0.28 | 0.29 |
| N | 1 | 0.109 | 1.555 | 1.5 | 1.485 | 1.425 | 1.4 | 1.385 | 1.411 |
|  | 5 | 0.097* | 1.041* | 1.01* | 0.999* | 0.443* | 0.802* | 0.785* | 0.825* |
|  | 10 | 0.082 | 0.963 | 0.721 | 0.476 | 0.125 | 0.576 | 0.426 | 0.346 |
| Mg | 1 | 0.032 | 0.028 | 0.035 | 0.043 | 0.038 | 0.040 | 0.03 | 0.038 |
|  | 5 | 0.03 | 0.024 | 0.03* | 0.036* | 0.033* | 0.033* | 0.03 | 0.032* |
|  | 10 | 0.025* | 0.017* | 0.022 | 0.025 | 0.028 | 0.027 | 0.024* | 0.025 |
| Ca | 1 | 20.1 | 25.22 | 20.46 | 23.5 | 20.15 | 21.12 | 22.6 | 22.46 |
|  | 5 | 19.3* | 23.35* | 17.5* | 21.75* | 18.92* | 19.8* | 21.2* | 19.45* |
|  | 10 | 17.45 | 19.87 | 14.15 | 18.67 | 14.74 | 15.98 | 18.64 | 15.4 |
| AVP | 1 | 2.5 | 3.1 | 2.75 | 2.65 | 2.6 | 3.0 | 2.9 | 2.7 |
|  | 5 | 2.3* | 3.1 | 2.65* | 2.1* | 2.15* | 2.65* | 2.7* | 2.55* |
|  | 10 | 1.97 | 2.75* | 2.42 | 1.85 | 1.85 | 2.15 | 2.33 | 2.15 |

a) Values = mean. (b)\*mean significant to others in the column of the same parameters at $p \leq 0.05$

**Table 3:** Values of TPH, Organic Carbon and Organic Matter in Various Soil Samples Treatment inside Seed Bags

| Parameter | Week | Samples |  |  |  |  |  |  |  |
| --- | --- | --- | --- | --- | --- | --- | --- | --- | --- |
|  |  | Control 1 | Control 2 | *P. aeruginosa* | *Flavobacterium* sp. | *Micrococcus* sp. | *Rhizopus* sp. | *Fusarium* sp | *Penicillium* sp. |
| Total Petroleum Hydrocarbons (PPM) | 0 | 2.41 | 20.29 | 20.29 | 20.29 | 20.29 | 20.29 | 20.29 | 20.29 |
|  | 3 | 2.36 | 19.21 | 13.25 | 13.02 | 15.36 | 16.19 | 9.28 | 16.46 |
|  | 6 | 2.28 | 19.01 | 10.2 | 10.42 | 12.46 | 13.12 | 5.75 | 9.56 |
|  | 10 | 2.09 | 18.78 | 2.19 | 3.18 | 4.1 | 4.12 | 2.25 | 3.87 |
| % Organic Carbon | 0 | 1.09 | 15.55 | 15.55 | 15.55 | 15.55 | 15.55 | 15.55 | 15.55 |
|  | 3 | 1.05 | 13.69 | 10.28 | 10.13 | 11.83 | 11.7 | 7.15 | 10.67 |
|  | 6 | 0.97 | 10.41 | 7.85 | 8.02 | 9.98 | 10.1 | 4.43 | 7.26 |
|  | 10 | 0.2* | 5.17* | 1.69* | 2.45* | 3.54* | 3.4* | 1.73* | 2.67* |
| % Organic Matter | 0 | 1.88 | 26.81 | 26.81 | 26.81 | 26.81 | 26.81 | 26.81 | 26.81 |
|  | 3 | 1.81 | 23.6 | 17.72 | 17.5 | 20.39 | 20.17 | 12.33 | 18.39 |
|  | 6 | 1.67 | 17.95 | 13.53 | 13.83 | 17.21 | 17.41 | 7.64 | 12.5 |
|  | 10 | 0.34* | 8.91* | 2.91* | 4.22* | 6.1* | 5.86* | 2.98* | 4.6* |

a.) Values = mean. (b)\*mean significant to others in the column of the same parameters at $p \leq 0.05$

Three (3) bacteria namely; *Pseudomonas* sp, *Flavobacterium* sp and *Micrococcus* sp and three fungal isolates (*Fusarium* sp, *Rhizopus* sp, and *Penicillium* sp) were isolated from the soil contaminated with spent engine oil using oil agar medium. Khan and Rizoi (2011); Abioye *et al.* (2012) and Jesubunmi (2014) also isolated similar bacteria in their works. Some of the fungi isolated from this study have been reported as hydrocarbons utilizers (April *et al.* (2000); Obire *et al.* (2008); George-Okafor *et al.* (2009) and Omotayo *et al.* (2012).Ayandele *et al.* (2012) also reported the biodegradation abilities of different genera of bacteria and fungi in their study. It has also been reported that anthropogenic impacts, such as changes in nutrients composition, have the potential to directly or indirectly affect the bacterial and fungal composition of the soil (Rousk *et al.*, 2009). Factorial Analysis of variance (ANOVA) was calculated for the physico-chemicals parameters, % OM, % OC and TPH and Heavy metals analysis of samples treated with different microbes as the weeks of treatment progressed. There was a significant difference among the parameters observed in this study and the activities of the microbes on the samples. The pH of all the samples was significantly different as the days progressed, except for the pH of the Control 2, which showed no significant difference at pH of 7.2 on week 1, 7.3 on week 5 and a pH of 7.4 on week 10. There was a significant difference in the potassium level, with the highest level recorded for Control 1 on week 1 (0.48) and followed by *Fusarium* sp. on week 1 with 0.45; while the lowest was recorded for control 2 on week 10 (0.23) followed by 0.27 recorded for *P. aeruginosa*. Okonokhua *et al.* (2007) reported that there was no significant difference between the pH of the control and spent oil treated soil in their study while Shahida *et al.* (2015) reported pH of neutral from 7.02-7.5 in different points and slightly alkaline (8.44) in one point of soil contaminated with hydrocarbons. Nitrogen showed a significant difference among the values with the activities of microorganisms as the weeks of treatment progressed, the highest value of 1.555 was recorded for Control 2 on week 1 and the lowest being 0.082 for the Control 1 on week 10. The lowest trend was found in the activities of *Micrococcus* sp. with the values ranging from 1.425 to 0.125 between Week 1 and Week 10. Significant difference was observed in the values of Mg and Ca, while there was a significant difference among the parameters tested during the experiments as Average values of Phosphorous (AVP) concentration from the highest (3.1) in the Control 2 was reduced to 1.85 by *Flavobacterium* sp and *Micrococcus* sp, which was the lowest value in week 10.

There was a reduction in total petroleum hydrocarbon concentrations in all the treatment with Control 1 having the lowest value of 2.09 followed by *P. aeruginosa* (2.19). Percentage organic carbon which was the highest (15.55) in Control 2 at the beginning of the experiment was actually low in control 1 from week 1 till the end of the experiment, while other than the result of control 1 were reducing gradually with *P. aeruginosa* having the lowest value on week 10 of the experiment. The percentage organic matter was low throughout in Control 1 with the upper limit being 1.88 and the lower being 0.34, while from the upper limit of 26.81for the other treatments, the parameter reduced gradually as the days of experiments progressed with the lowest value being 2.91 with *P. aeruginosa*. There was a significant difference on week 10 compared with the other weeks and also among other treatments.

There was a significant difference in the values of Fe, Zn, Pb, Cd, Cu, Cr and Ni during the experimental period with control 1 having the lowest values on week 10 for Zn (0.11), Pb (0.02) and Cd (0.026) while both the control 1 and *Rhizopus* sp. has the lowest value for Fe, control 2 has the lowest value of Cr (0.06) but *Fusarium* sp. has the lowest for Cu (0.11) and Ni (0.07) on week 10 of the treatment. *Flavobacterium* sp. has the lowest value of Cr after treatment when they were compared to the initial values at the beginning of the experiments (Table 4).

**Table 4:** Heavy Metal Analysis of the Treatments during the Experimental Period

| Parameter | Week | Samples |  |  |  |  |  |  |  |
| --- | --- | --- | --- | --- | --- | --- | --- | --- | --- |
|  |  | Control 1 | Control 2 | *P. aeruginosa* | *Flavobacterium* sp. | *Micrococcus* sp. | *Fusarium* sp. | *Rhizopus* sp | *Penicillium* sp. |
| Fe | 1 | 0.016* | 0.096 | 0.026 | 0.036 | 0.025 | 0.021 | 0.018 | 0.019 |
|  | 5 | 0.012 | 0.07* | 0.021 | 0.029* | 0.018* | 0.017* | 0.014 | 0.015* |
|  | 10 | 0.009 | 0.045 | 0.015* | 0.018 | 0.01 | 0.011 | 0.009* | 0.011 |
| Zn | 1 | 0.28 | 0.45 | 0.3 | 0.33 | 0.3 | 0.35 | 0.38 | 0.37 |
|  | 5 | 0.17* | 0.39* | 0.25 | 0.28* | 0.25 | 0.29* | 0.3* | 0.31* |
|  | 10 | 0.11 | 0.27 | 0.19* | 0.21 | 0.19* | 0.2 | 0.22 | 0.26 |
| Pb | 1 | 0.04 | 0.35* | 0.28* | 0.31 | 0.34* | 0.32 | 0.32 | 0.36 |
|  | 5 | 0.03 | 0.38 | 0.31 | 0.32 | 0.36 | 0.31 | 0.29* | 0.34* |
|  | 10 | 0.02 | 0.39 | 0.33 | 0.34* | 0.37 | 0.29* | 0.26 | 0.29 |
| Cd | 1 | 0.09* | 0.15 | 0.095 | 0.097 | 0.095 | 0.09 | 0.09 | 0.092 |
|  | 5 | 0.045 | 0.13 | 0.054* | 0.07* | 0.065* | 0.06* | 0.08 | 0.079 |
|  | 10 | 0.026 | 0.1 | 0.031 | 0.043 | 0.031 | 0.035 | 0.065* | 0.054* |
| Cu | 1 | 0.17 | 0.26 | 0.21* | 0.2 | 0.23 | 0.22 | 0.21 | 0.22 |
|  | 5 | 0.15* | 0.24* | 0.17 | 0.17* | 0.2 | 0.19 | 0.19 | 0.18 |
|  | 10 | 0.13 | 0.21 | 0.12 | 0.13 | 0.16* | 0.11* | 0.16* | 0.13* |
| Cr | 1 | 0.15 | 0.25 | 0.17 | 0.15 | 0.16 | 0.14 | 0.19 | 0.15 |
|  | 5 | 0.14 | 0.23* | 0.13* | 0.11* | 0.12* | 0.11* | 0.17* | 0.13* |
|  | 10 | 0.11* | 0.19 | 0.08 | 0.06 | 0.07 | 0.07 | 0.11 | 0.09 |
| Ni | 1 | 0.15 | 0.25 | 0.17 | 0.15 | 0.16 | 0.14 | 0.19 | 0.18 |
|  | 5 | 0.14 | 0.21* | 0.13* | 0.11* | 0.13* | 0.11* | 0.15* | 0.15* |
|  | 10 | 0.13* | 0.19 | 0.08 | 0.08 | 0.09 | 0.07 | 0.11 | 0.1 |

a.) Values = mean. (b)\*mean significant to others in the column of the same parameters at $p \leq 0.05$

There have been different reports on the effects of hydrocarbons on the physicochemical properties, TPH, OM, OC and heavy metals analysis of contaminated soils. pH levels of the soil samples increased from acidic to neutral in all the soil samples, Controls 1 and 2 and all the treated soil sample. pH of 6.62-7.72, phosphate of 0.10-0.11mg/kg, Nitrate of 0.22-0.31mg/kg, %TOC of 1.23-3.99%, Potassium of 0.87-0.90mg/kg and Nitrogen of 0.06-0.51% have been reported by Omotayo *et al.* (2012) in their study of crude oil degradation by microorganisms in soil composts. Shahida *et al.* (2015) also observed the high content of potassium and sodium contents from all the soil samples analysed in their work and low contents of available carbon, nitrogen, and phosphorus. While Adenipekun *et al.* (2013) reported increased in the level of % OC, % OM, Phosphorus and Potassium after 2 months of biodegradation of soil contaminated with spent and fresh cutting fluids and decrease in total nitrogen and pH. The increase in the percentage of organic carbon and nitrogen of spent engine oil soil in relative to control could be attributed to the effect of spent engine oil (Okonokhua *et al.*, 2007) while Kayode et al. (2009) also reported a reduction in nitrogen content of their soil treated with spent lubricant oil. While available carbon, nitrogen, and phosphorus were very low in the study of Shahida et al. (2015), Lovely and Cackette (2001) observed significant increase in the nitrogen and carbon contents in their own study and reported that increase in nitrogen might be due to the increased in atmospheric nitrogen during the degradation process, while carbon increase might be due to the presence of carbons in hydrocarbons.

Atlas, (1984) reported that nutrient availability is dependent on the nature of the environment sometimes and that nutrients such as nitrogen and phosphorus could be limiting sometimes and thus affecting the biodegradation process. Also, the presence of high numbers of hydrocarbon degraders has been linked to the high availability of nutrients (Atlas, 1995; Ijah and Okang, 1993).

Reduction in Total Petroleum Hydrocarbons (TPH) continues to increase as the weeks of biodegradation increased, the highest TPH removal was observed in *P. aeruginosa* (2.19 PPM), followed by *Fusarium* sp (2.25) while soil treated with *Rhizopus* sp had the highest TPH content (4.12) at the end of the experiment (Table 3). Adenipekun and Isikhuemhen (2008) had reported that TPH removal always increases as the days of incubation increases, while Shukor *et al.* (2009) reported that biodegradation rate of *Pseudomonas* sp in diesel oil contaminated soil increased as the days of biodegradation increased. Abdulsalam *et al.* (2012) also reported an increase in removal of TPH in soil contaminated with used motor oil increased as the incubation period is elongated, while Adenipekun *et al.* (2013) reported TPH decreases after two (2) months from fresh and spent cutting oil contaminated soil samples.

There was a gradual and significant reduction in the content of heavy metals present in the soil contaminated with spent engine oil in relative to the control soils (table 4). Adenipekun *et al.* (2013) also observed decreased in Mn, Pb, Ni, and Cu in soil contaminated with fresh and spent cutting fluid after 2 months even in the control. While Adams *et al.* (2014) reported a gradual decrease in the different metals analysed in the spent oil contaminated site as the experimental days progressed.

## CONCLUSION

The results obtained from this study showed that all organisms isolated from the contaminated soil can be used for biodegradation process. Reduction in total petroleum hydrocarbons and heavy metals present in the contaminated soil samples were observed when all the isolated organisms were used in the treatment of the contaminated soil samples.

